# Ratatoskr: A tool for automated retrieval of taxonomic type strain sequences and metadata

**DOI:** 10.64898/2026.01.26.700362

**Authors:** Christopher Turkington, Fabian Bastiaanssen, Neda Nezam-Abadi, Andrey Shkoporov, Colin Hill

## Abstract

Bacterial taxonomic type strains anchor species names to physical and genomic reference material, making them essential for reproducible and comparable prokaryotic research. While reference strains are often well-characterised through curated metadata, nomenclature histories, and sequence records, no single database holds up-to-date information on all these aspects, resulting in fragmented information. Gathering the complete set of information for a type strain is further complicated by inconsistencies in nomenclature between sources due to the often-numerous synonyms that can describe a single strain. As a result, collecting type strain data for taxonomic proposals and emendations can be an onerous task requiring extensive manual curation. To address this issue, we introduce Ratatoskr, a Python-based tool that automates the retrieval of sequences and metadata for bacterial taxonomic type strains. Ratatoskr facilitates this by collecting the latest type strain information of the List of Prokaryotic names with Standing in Nomenclature (LPSN) and using this information to query the BacDive and NCBI databases. By applying known taxonomic synonym information Ratatoskr is able to resolve cross-database inconsistencies and streamline the retrieval process. We show that through its use, Ratatoskr can obtain metadata and sequence data for type strains of bacteria within minutes to seconds, depending on the number of members within the requested taxon. By automating this retrieval, Ratatoskr provides fast, accurate, and readily shareable starting points for studies involving the use of taxonomic type strains and data, such as new taxonomic proposals or emendations.

**Data summary:** Ratatoskr was developed using Python 3 and is freely available at https://github.com/Fabian-Bastiaanssen/Ratatoskr under a GPL-3.0 licence.

## Introduction

The proposal, classification, and revision of bacterial taxonomic boundaries are essential aspects of prokaryotic research [1, 2]. Central to this effort are taxonomic type strains that serve as reference points for the physical and genomic characteristics of a taxon, forming the basis for accurate and reproducible delineation [3]. It is therefore essential that the scientific community has access to accurate and up-to-date information on these organisms to allow taxonomic analyses to be conducted.

The List of Prokaryotic names with Standing in Nomenclature (LPSN) was created as a central repository for taxonomic names and synonyms recognised under the rules of the International Code of Nomenclature of Prokaryotes (ICNP) [4, 5]. Today, the LPSN is actively curated and maintained as part of the DSMZ Digital Diversity database collection [6], and is the primary reference resource for information on taxonomic type strains. However, while the LPSN provides current and extensive information on the nomenclature and revision history (including homotypic synonyms, proposal references, and some 16S rRNA sequences) it does not contain information on the phenotypic characteristics or genomic sequences of these organisms; information that is typically reported or compared within taxonomic proposals and emendations [7, 8].

Experimentally determined phenotypic characteristics of taxonomic type strains (as well as other non-type isolates) can instead be found on BacDive – another database curated and maintained by the DSMZ [9]. BacDive represents the largest database of standardised information on prokaryotic phenotypic research data, including information on ~98% of validly described bacterial species [10]. BacDive also contains some accession numbers to genome and 16S rRNA sequence data in external sequence repositories, such as NCBI GenBank [11], JGI IMG [12] and BV-BRC/PATRIC [13]. However, this information is not necessarily available for all type strains, including those instances where genome/16S sequences do exist within public sequence databases.

The most direct source for taxonomic type isolate genome and 16S rRNA sequences are the collections of the International Nucleotide Sequence Database Collaboration (INSDC), namely GenBank, the European Nucleotide Archive (ENA), and the DNA Databank of Japan (DDBJ) [14]. However, retrieving sequence information on taxonomic types from these databases is not necessarily straightforward as entries do not always directly follow the current ‘correct’ nomenclature. Instead, entries follow a distinct taxonomy that can result in taxonomic types being listed in these databases under their alternative binomial synonyms [15]. While taxonomically acceptable (as these binomials are valid synonyms) this can complicate search efforts by resulting in naming conflicts between databases [16], a problem that becomes more pronounced when strain-level synonyms are also incorporated.

As no single database integrates nomenclature, phenotypic characteristics, and nucleotide sequences, initiation of studies proposing new taxa or revising existing descriptions can become a laborious and protracted undertaking, especially when addressing higher-rank taxa or groups with extensive membership. Here, we introduce Ratatoskr, a Python-based tool designed to streamline the retrieval of taxonomic type strain sequence data and metadata. Ratatoskr does this by providing a single-entry point to retrieve and unify the plethora of data available on the LPSN, BacDive, and NCBI databases describing the users taxon of interest. By accessing these databases with each user call and providing a structured set of outputs, Ratatoskr ensures the generation of up-to-date, consistent, and easily shareable datasets well-suited for downstream analysis within open research frameworks.

## Methods

### Overview of the Ratatoskr pipeline

#### Inputs

Ratatoskr only requires three sets of inputs from the user. Firstly, the user must provide the taxonomic name they wish to retrieve information for (e.g. *Pasteurellaceae*) along with the desired path to where the output folder should be created. Any taxonomic level above species can be provided as input but it is important that this taxonomic name is the current taxonomic ‘correct name’ for the taxon of interest. The current correct name for a taxon can be checked/obtained by the user by using their query as a search term on the LPSN website (https://lpsn.dsmz.de/). During the run, the user will then be prompted to provide their DSMZ credentials (email and password) for accessing the LPSN API, and their NCBI API key to retrieve data through NCBI Datasets. Users can create an account through the LPSN API webpage (https://api.lpsn.dsmz.de/), and NCBI API keys can be generated via the NCBI API webpage (https://www.ncbi.nlm.nih.gov/datasets/docs/v2/api/).

#### Type strain information and sequence retrieval

Initially, all type strain information in the LPSN database is retrieved using the LPSN API tool (https://github.com/LeibnizDSMZ/lpsn-api; [17]), specifically a modified version of this tool that allows asynchronous retrieval of returned pages (https://pypi.org/project/async-dsmz/), increasing retrieval speed when large numbers of results are returned. This dataset is then converted into a parsable data frame using Polars (https://pola.rs/) which is then searched for the user requested taxon and all associated entries retrieved. Along with taxon membership, additional information held in LPSN that are useful for later steps in the Ratatoskr pipeline are also collected at this stage (e.g. 16S rRNA accessions, and all binomial and strain name synonyms), together with other general metadata that may be of interest to users (e.g. the references for species proposal and listing, their associated authority, and their LPSN ID’s).

All collected taxa are then used to query the BacDive database using the BacDive API tool (https://github.com/JKoblitz/bacdive-api; [10]), again using a modified version of this tool for asynchronous retrieval of results (https://pypi.org/project/async-dsmz/). During this step, phenotypic data for the taxonomic types are collected where available (e.g. culture conditions, morphological data, Analytical Profile Index (API)-associated results, and metabolite utilisation/production data), as well as any 16S rRNA accessions not found during the LPSN search, plus any genome accessions. Where multiple possible genome sequences are available, preference is given to genomes with higher levels of assembly completion. For 16S rRNA sequences, if multiple accessions are present the returned accession will be that of the longest length sequence associated with the entry (up to a maximum of 2 kb). In addition, any further synonyms listed in BacDive not retrieved from the LPSN search are also collected at this stage.

For each type strain NCBI taxon ID’s are then collected using the Entrez module from the Biopython library using all synonyms obtained from the previous steps as queries [18]. For types with no 16S rRNA sequence from the LPSN or BacDive searches, these taxon IDs (plus any binomial synonyms associated with them) are then used to identify the missing 16S rRNA sequences via Entrez. Similarly, for those with no genome accession at this stage, NCBI taxon IDs and binomial synonyms are used to search for genome accessions in NCBI using NCBI Datasets (https://github.com/ncbi/datasets; [19]). After these steps, 16S rRNA and genome accessions collected at these and the previous steps are used to retrieve the associated sequences through Entrez and NCBI Datasets, respectively.

#### Outputs

Ratatoskr generates three primary sets of outputs: taxonomic information, sequence data, and characteristic metadata. For taxonomic information, a folder is produced containing a tab-separated file that includes descriptions of each entries associated taxonomic levels, reference for effective publication, and authority, among other details. For characteristic metadata, a folder is generated that contains three tab-separated files and a subfolder. The three files represent one of the following sets of characteristics available on BacDive: fatty acid profiles, metabolite utilisation, and phenotypic metadata (e.g. growth conditions, morphology, compound production etc). In the subfolder are tab-separated files containing the results for each API strip associated with the type strains (one file per API strip). For sequence data a folder is created that holds one file and two subfolders. The file includes metadata for each type strain’s retrieved sequences (genome and 16S rRNA accessions), and each subfolder contains either all 16S rRNA sequences in FASTA format or the genome sequences, which are further divided into two subfolders: one with genomes in FASTA format and another with genomes in GenBank format.

### Retrieval Benchmarking

To create a ground truth dataset for benchmarking type strain retrieval, we based our test dataset on the type strains listed as nomenclature correct within the *genera, species and subspecies* (GSS) list (2025-12-15 version) available from the LPSN download page (https://lpsn.dsmz.de/downloads), which contains bulk nomenclature information for taxon from the genus-level and below. Here, we only included those taxa for which a full taxonomic linage could be retrieved using the LPSN API (accessed via async-dsmz v2025.0.4; https://github.com/Fabian-Bastiaanssen/async_dsmz). Using this method, we were able to assign all but 15 of the 22,037 correct name entries in the GSS to their higher-level classifications. Manual examination of these fifteen outliers identified these as orphan correct names; i.e. correct names where their associated parent taxon is no longer described as taxonomically correct. As these fifteen represented orphan taxa, we excluded these from our taxonomic reference dataset. The remaining 22,022 taxonomic types listed in GSS were then used as the ground truth dataset for benchmarking.

We compared the ability of Ratatoskr in retrieving type strain taxonomic lineages, 16S rRNA sequences, and genome sequence data to each of the individual APIs it calls from, namely LPSN, BacDive, and NCBI APIs (accessed as of 15th December 2025). For the LPSN and

BacDive APIs, genera, species and subspecies names in the GSS were used as inputs to query their RESTful APIs with async-dsmz v2025.0.4 (https://github.com/Fabian-Bastiaanssen/async_dsmz). Correct strain matches were then determined based on strain-name synonyms given in the GSS. Full taxonomy given for the produced hit(s) were then compared to that obtained from the LPSN for the corresponding GSS entry and matches/divergencies from this taxonomy then collated. At this point, any 16S rRNA or genome accession provided for the entry was also collected. For NCBI queries, NCBI taxonomic IDs were collected for all species and subspecies entries using the taxonomy endpoint of the NCBI Datasets v2 RESTful API. The resulting NCBI taxon IDs were used to query genomes via the NCBI Datasets API, and corresponding 16S rRNA sequences were retrieved using Biopython v1.86 as interface to the E-utilities nucleotide endpoint. For all methods, any missing entries were marked as taxonomically incorrect at all levels, while successful 16S rRNA/genome accession retrieval was defined as collection of a sequence matching any of the strain name synonyms associated with the type isolates given in the GSS list.

Retrieval times.were also examined by running all possible taxonomic queries from genus to domain through the Ratatoskr pipeline. These tests were carried out with the flag *‘--skip_download’* to isolate the influence of internet speeds and file sizes, thus these tests do not include the download of 16S rRNA or genome sequences in their runtime. However, we do present examples of runs conducted for a subset of fifty distinct taxonomic queries – covering ten randomly selected representatives from phylum, class, order, family, and genus – to illustrate estimated overall runtimes inclusive of sequence download.

The above benchmarking was performed using Python v3.12, and all graphs were created using R version 4.3.1, patchwork v1.2.0 (https://patchwork.data-imaginist.com/), tidyverse v2.0.0 [20], and ggh4x v0.3.0 (https://github.com/teunbrand/ggh4x). All source data and code is archived on Zenodo (https://doi.org/10.5281/zenodo.18327275).

## Results and Discussion

To determine the breadth and accuracy of taxonomic lineage, 16S rRNA, and genome sequence retrieval through Ratatoskr, we examined its ability to collect this information for each of the 22,022 correct species/subspecies entries within the LPSN GSS list at time of writing (see methods).

First, we examined retrieval of taxonomic data using either solely the LPSN, BacDive, and NCBI APIs, or Ratatoskr (Figure 1). These APIs were selected as these APIs represent means of accessing three databases that hold taxonomic and sequence information and also are accessed within the Ratatoskr pipeline. From this, it was observed that querying the BacDive and NCBI APIs led to the return of examples of inaccurate taxonomic information at each level. This was particularly true for the BacDive database at the kingdom and phylum levels where > 50% of taxa information returned was incorrect. For kingdom, this is explained through the systematic absence of kingdom level taxonomic information in the BacDive database. However, for phylum and all other instances conflicts were the result of either sporadic missing or conflicting information being returned instead of the correct nomenclature. This, is interesting as although BacDive is said to be directly updated with reference to the LPSN taxonomy [10], it performed worse than NCBI at every level except the strain level. NCBI in contrast acknowledges that its taxonomy can deviate from that of the current nomenclature [15]. The LPSN API identified all current nomenclature-correct GSS members at all taxonomic levels because the LPSN originated the GSS list used in the benchmarking. Likewise then, as Ratatoskr uses the LPSN API within its pipeline to retrieve taxonomic information it also retrieved all correct taxonomic information.

**Figure 1.**
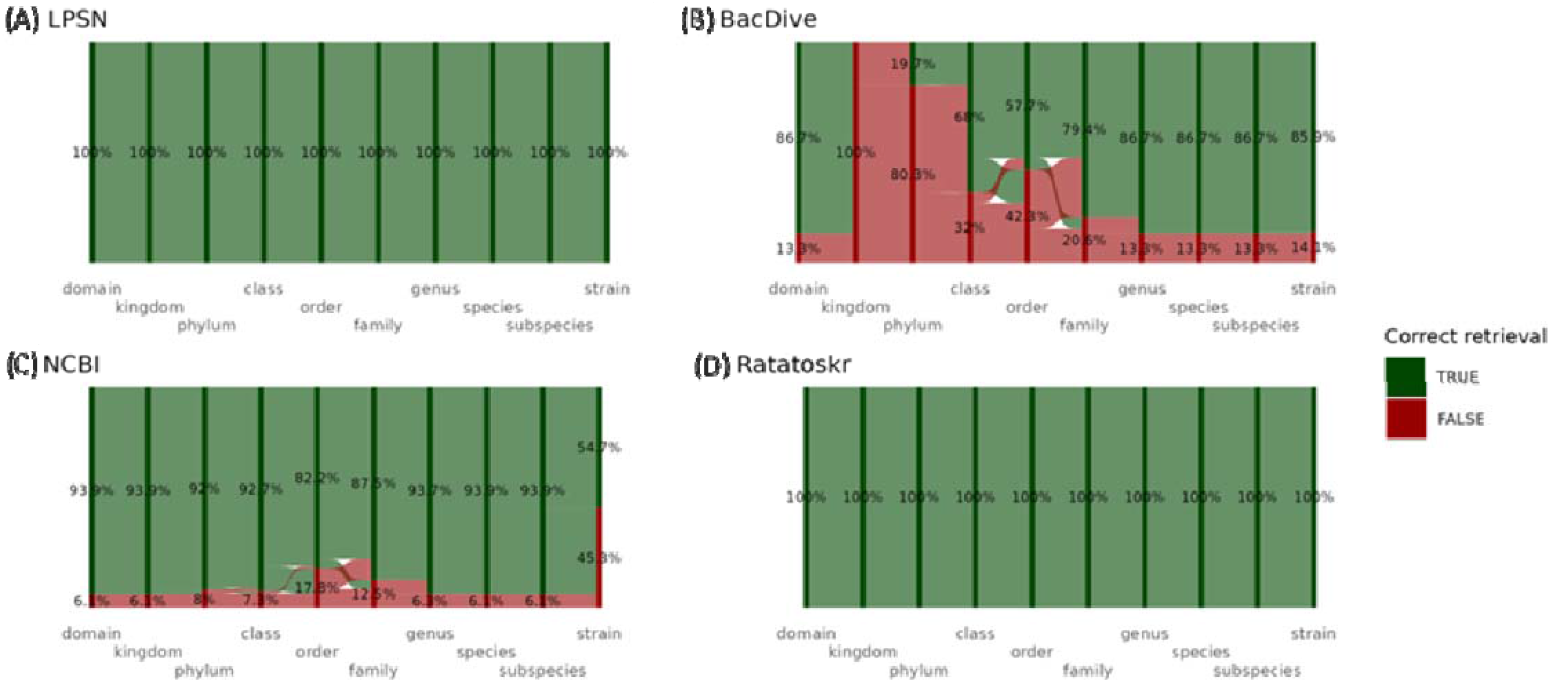
Sankey plots showing the ability of API retrieval methods to collect the correct nomenclature type strains for each taxonomic level, namely the (A) LPSN API, (B) BacDive API, (C) NCBI API, plus (D) Ratatoskr. Each horizontal stage represent a taxonomic level, with the percentage value and size of the coloured area corresponding to the proportion of strains placed at that taxonomic level. Green nodes indicate correct nomenclature matches, and red nodes indicate incorrect retrievals (i.e. either discrepancies or entirely missing entries). The flow between stages indicates changes in retrieval accuracy from one stage to the next.

We next examined the ability of Ratatoskr and the API tools to retrieve 16S rRNA and genome sequence data for each of the type entries within the GSS list (Figure 2). For 16S rRNA sequences, LPSN was able to recover an accession for 96.5% of type strains, considerably more than ∼75% and ∼55% retrieved using the BacDive and NCBI APIs, respectively. In the case of BacDive, most of the unsuccessful retrievals were due to entries containing no accession information and thus not being returned, meanwhile for NCBI, the unsuccessful entries were actually caused by returning incorrect strain/taxon. This occurred when type strain sequences were linked to entries with incorrect taxonomic IDs, and NCBI instead returned sequences from different strains of the species (e.g. the 16S rRNA accession U65012 for *Stutzerimonas stutzeri* ATCC 14405 was returned instead of the correct nomenclature type strain ATCC 17588). Ratatoskr was able to recover the greatest proportion of 16S rRNA accessions, collecting accessions for 98.7% of the GSS type strains. This increase was due to its retrieval of an additional 2.2% of accessions through the BacDive and NCBI databases that were not available through the LPSN entry.

**Figure 2.**
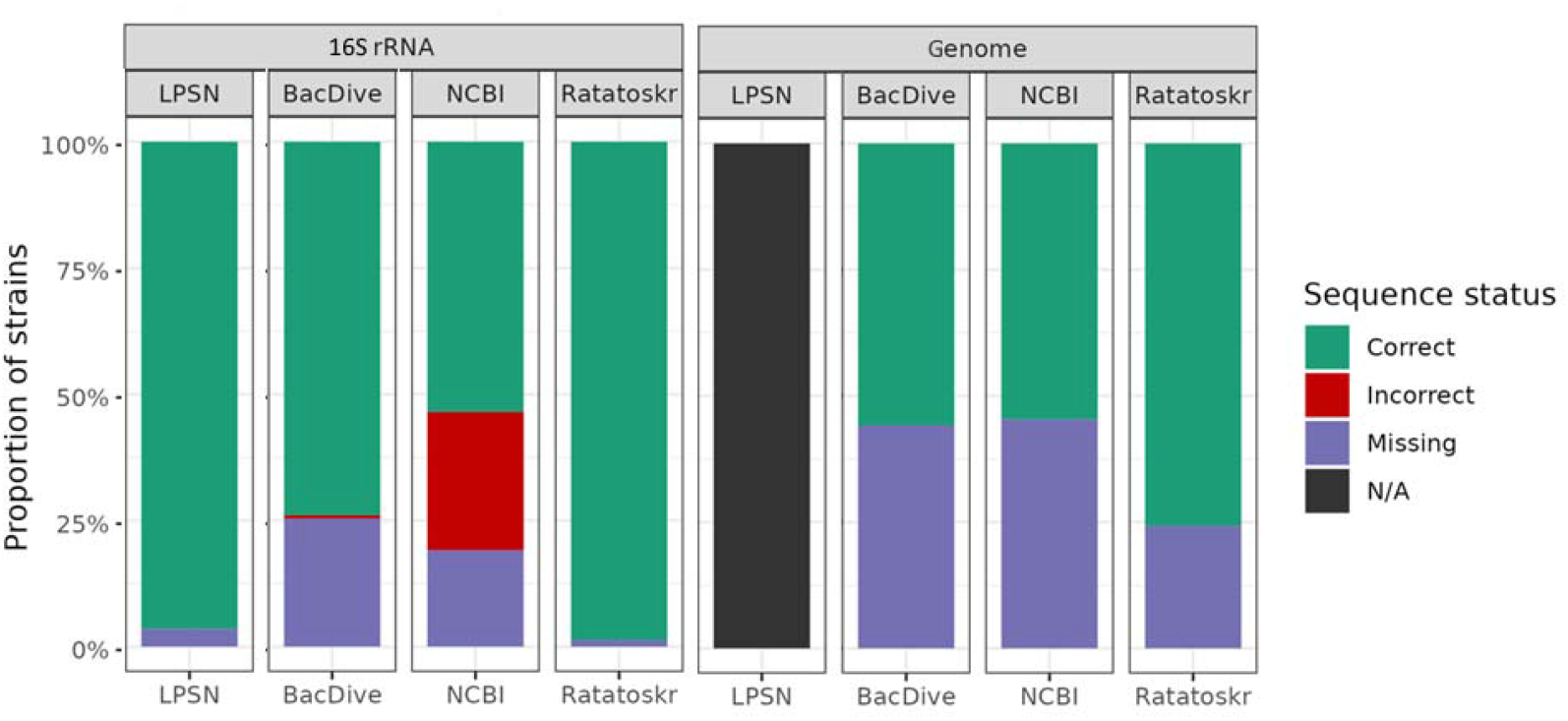
Genome and 16S rRNA retrieval results using species and strain queries from the LPSN GSS list using either the LPSN API, BacDive API, NCBI API, or Ratatoskr. Each section in a bar displays the proportions of queries that returned a sequence matching the target strain (green sections), an incorrect sequence (i.e. matching a different strain of the correct species; red sections), or where no result was returned (purple section). Note that LPSN does not hold genome sequence data or accessions at all and is therefore labelled as N/A in the genome sequence plot (black). Here, queries described as missing/incorrect results do not mean a correct sequence for that strain does not exist within the database, but rather that querying the correct taxon and its strain name synonyms did not lead to its return.

For genome data, while the LPSN database lacks genome sequence data altogether (thus could not include for comparison), the BacDive and NCBI APIs were able to gather accessions for > 50% of the GSS type strains in comparable numbers (55.8 and 54.7%, respectively). In both their cases the unsuccessful searches were due to queries not returning any entries rather than the wrong query. Ratatoskr was able to retrieve a greater number of accessions than either of these APIs alone, recovering a total of 75.7% of members from the GSS list. This is again due to Ratatoskr’s dual access to each of these databases, which allows the information they contain to complement one another and thereby increase the number of entries retrieved. However, missing results were observed for this method too, indicating either a genuine absence of strain-specific sequences or that available entries are annotated in a way that none of the methods used here, nor the composite search Ratatoskr employs, can retrieve. Ratatoskr’s success in retrieving a greater number of accessions for both 16S rRNA and genome sequences than any of the APIs alone is likely also driven by its use of binomial and strain synonyms that allows it to pick up entries missed by each of the individual APIs when only the nomenclature correct names are used.

Finally, we also examined the runtime for deployment of Ratatoskr to recover information across various taxonomic levels. In the first set of runtime tests, we examined the time required for Ratatoskr to retrieve each currently recognised nomenclature-correct taxon (based on the full taxonomy obtained for entries in the GSS list). Here, unique names at each taxonomic rank were used as input to Ratatoskr and runtimes obtained in the absence of sequence download (this was done to remove the influence of local download speed and file size in entry collection). As might be expected, runtimes were found to increase proportionally with the number of members per query rather than taxonomic rank (Figure 3A). However, even for the largest query – recovering the entirety of type strain sequence accessions and metadata for the domain bacteria – Ratatoskr was able to return data in around 25 minutes, a job that without automation would likely take days. Further examination of runtimes, this time including sequence file download in the run, using a subset of fifty of these taxon (ten randomly selected examples each for the ranks of phylum, class, order, family, and genus), showed that inclusion of this step leads to runtimes being roughly tripled (∼2.7x) (Figure 3B). Thus this suggests that this step is the greatest time bottleneck to Ratatoskr’s completion. Regardless, these results demonstrate that Ratatoskr can complete the retrieval of taxonomy, metadata, and sequence data within a timeframe far faster than could be realistically achievable by manual curation.

**Figure 3.**
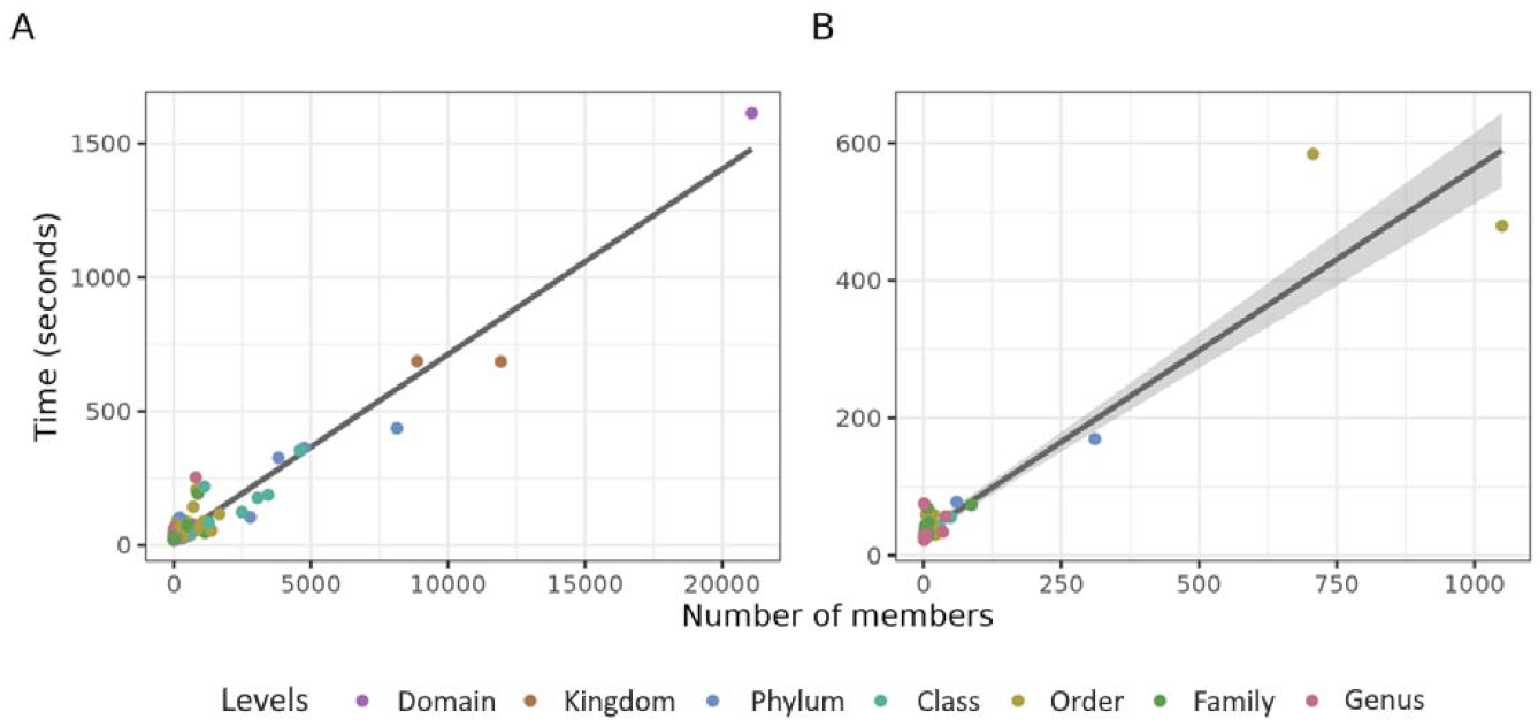
Observed Ratatoskr runtimes against the number of members returned per taxonomic query. (A) Illustrates Ratatoskr runtimes for all possible correct taxonomic queries in the nomenclature, excluding the steps for downloading 16S rRNA and genome sequences (i.e. taxonomy, metadata, and accession retrieval but no sequence download). (B) Runtimes for a subset of 50 queries (10 queries each at phylum, class, order, family, and genus levels) that included the sequence download steps. Each point represents a single input query, coloured by its taxonomic rank (e.g. Lachnospiraceae and *Bacteroides* would be coloured green and red, respectively). The trend line represents the best fit across the datasets with shaded regions indicating 95% confidence intervals.

### Limitations

Ratatoskr does not currently incorporate *Candidatus* taxa, something that could be problematic in some instances given the recent inclusion of the new Section 10 of the ICNP [21, 22]. *Candidatus* taxa (where proposed in-line with the criteria given in Section 10 of the ICNP) are now recognised as pro-valid and as such, proposal of novel taxon should examine if their proposed taxon corresponds to a previously described *Candidatus* taxon as to conform to the requirements of rule 72 [21]. Ratatoskr is currently unable to include these organisms because, although these organisms are listed on the LPSN website, they are not yet accessible through the LPSN API. However, should these entries become retrievable through the API in the future, they will be incorporated into a subsequent update of Ratatoskr.

One further point of note is that the coverage and accuracy of Ratatoskr necessarily depends on the robustness of the external databases it interrogates, and even well-maintained resources such as LPSN contain occasional errors. For example, at the time of writing, LPSN lists NCTC 14570 as a potential type-strain identifier for both *Luteimonas colneyensis* and *Microbacterium commune*, despite these species sharing only the rank of Bacteria. Both of these taxa were validly published as part of a large set of novel organisms isolated from the chicken gut [23, 24], with NCTC 14570 corresponding to strain Sa2BVA3, the type strain of *L. colneyensis* [24]. In contrast, the type strain of *M. commune* is Re1 and is referenced as being deposited under NCTC 14561 [24], an accession that does not appear in the NCTC catalogue (https://www.culturecollections.org.uk/). Although Ratatoskr would return the erroneous *M. commune* strain-name synonym from LPSN in its output, its workflow mitigates downstream influence of the error when accessing the BacDive and NCBI databases, as queries are initiated using the binomial name, and strain synonyms are then used only as filters on returned results. In this case, the incorrect synonym is a unique culture-collection number, preventing accidental overlap across taxa. However, if the error involved a more generic strain designation, there would be greater potential for incorrect data to be inadvertently returned. Thus, while such instances are rare, it is still recommended that the user still validates the retrieved data using their own knowledge of the taxa being studied.

## Conclusion

Overall, through its use Ratatoskr can prevent the need for researchers to undertake time-consuming manual curation of type strain sequence and metadata. In addition, the workflow and output implemented by Ratatoskr directly supports the FAIR (Findable, Accessible, Interoperable, Reusable) data principles that increasingly govern microbial research [25]. Ratatoskr enhances findability by consolidating disparate identifiers and synonyms into structured, queryable outputs. It improves accessibility through automated retrieval of openly available data from major public repositories. Its use of standard file formats in its output (TSV, FASTA, and GenBank) promotes interoperability, and by generating readily shareable datasets, Ratatoskr strengthens reusability, ensuring that taxonomic proposals can be easily replicated in future reanalyses. This allows Ratatoskr to fill a critical gap in microbial taxonomy by providing a robust, scalable, and FAIR-aligned approach for retrieving the metadata and sequence resources required for taxonomy-focused microbial studies.

## Author Contributions

CT: Conceptualization, Formal analysis, Investigation, Methodology, Software, Supervision, Validation, Writing - Original Draft, Writing - Review & Editing

FB: Conceptualization, Data Curation, Formal analysis, Investigation, Methodology, Software, Validation, Visualisation, Writing - Review & Editing, Writing - Original Draft

NNA: Project administration, Writing - Review & Editing AS: Resources, Supervision, Writing - Review & Editing

CH: Funding acquisition, Resources, Supervision, Writing - Review & Editing

Conflict of interest

The authors declare no conflicts of interest.

## Acknowledgements

The authors would like to thank the International Committee on Systematics of Prokaryotes (ICSP), INSDC databases, and those at DSMZ, as without their work keeping taxonomic information, characteristics, and sequencing data up-to-date and well maintained, such work would not be possible.

